# Characterization of a gene-trap knockout mouse model of *Scn2a* encoding voltage-gated sodium channel Nav1.2

**DOI:** 10.1101/2020.06.23.150367

**Authors:** Muriel Eaton, Jingliang Zhang, Zhixiong Ma, Anthony C. Park, Emma Lietzke, Chloé Maricela Romero, Yushuang Liu, Emily Rose Coleman, Xiaoling Chen, Tiange Xiao, Zhuo Huang, William C. Skarnes, Wendy A. Koss, Yang Yang

## Abstract

Recent large-scale genomic studies have revealed *SCN2A* as one of the most frequently mutated gene in patients with neurodevelopmental disorders including autism spectrum disorder and intellectual disability. *SCN2A* encodes for voltage-gated sodium channel isoform 1.2 (Nav1.2), which is mainly expressed in the central nervous system and responsible for the propagation of neuronal action potentials. Homozygous knockout (null) of *Scn2a* is perinatal lethal, whereas heterozygous knockout of *Scn2a* results in mild behavior abnormalities. To achieve a more substantial, but not complete, reduction of *Scn2a* expression, we characterized a *Scn2a* deficient mouse model using a targeted gene trap knockout (gtKO) strategy to recapitulate loss-of-function *SCN2A* disorders. This model produces viable homozygous mice (*Scn2a*^*gtKO/gtKO*^) that can survive to adulthood, with markedly low but detectable Nav1.2 expression. Although *Scn2a*^*gtKO/gtKO*^ adult mice possess normal olfactory, taste, hearing, and mechanical sensitivity, they have decreased thermal and cold tolerance. Innate behaviors are profoundly impaired including impaired nesting, marble burying, and mating. These mice also have increased food and water intake with subsequent increases in fecal excretion of more but smaller fecal boli. This novel *Scn2a* gene trap knockout mouse thus provides a unique model to study pathophysiology associated with *Scn2a* deficiency.

## INTRODUCTION

*SCN2A* encodes for the alpha subunit of the voltage-gated sodium channel isoform (Nav1.2), which is a major sodium channel expressed in the neurons of the central nervous system responsible for the initiation, propagation, and backpropagation of action potentials. During development (first through second year in humans or first through third week in mice), Nav1.2 is the main sodium channel isoform expressed in the axon of neurons, thereby making it the dominant initiator and propagator of action potentials in the central nervous system [1].

Mutation variants in *SCN2A* are strongly associated with neurodevelopmental disorders such as autism spectrum disorder (ASD), intellectual disability, epilepsy encephalopathy, and also schizophrenia with comorbidities including sleep irregularities, gastrointestinal disturbances, and motor discoordination [2] [3] [4] [5] [6]. An estimated 400 cases of *SCN2A*-related disorders are born each year in the United States [7]; unfortunately, there is no treatment available that is specific for *SCN2A*. A majority of *SCN2A* mutations identified from ASD cases are protein-truncating variants or have been experimentally verified to result in loss-of-function (*SCN2A* deficiency) [8]. Development of effective therapeutic interventions for patients carrying *SCN2A* loss-of-function variations will rely on a deeper understanding of cellular, circuital, and behavioral consequences of *SCN2A* deficiency.

To study disorders associated with loss-of-function *SCN2A* variants, *Scn2a* knockout mouse models were generated [9]. Canonical knock-out (null) of *Scn2a* in mice is perinatal lethal, therefore current studies focus on heterozygous *Scn2a* knockout mice (*Scn2a*^*+/*−^) as the main model for study, which display only mild behavioral abnormalities. *Scn2a*^*+/*−^ mice show little differences from wild-type (WT) mice in body weight, olfactory, auditory startle, thermal sensitivity, nesting, and marble burying [10] [11] [12] [13]. We reasoned that if we can substantially reduce the *Scn2a* expression level without eliminating it completely, the residual *Scn2a* protein may allow the mice to survive and to exhibit more severe phenotypes in adulthood than is observed in heterozygous knockout mice, thereby providing a novel model for the study of *Scn2a* deficiency-related biological processes. To this end, we have characterized a *Scn2a* deficient mouse model using a gene-trap knockout (gtKO) approach.

The gene-trap knockout is a versatile approach and is achieved by inserting a “trapping cassette” into an intron of the gene of interest [14] [15], thus trapping the upstream exons by splicing to the cassette and truncating the mRNA (**Figure 1A**). The trapping cassette contains a LacZ reporter flanked by a splice acceptor and polyadenylation signal, as well as site-specific recombination sites for further genetic modification of the allele. Mouse strains with a trapping cassette are designated as tm1a, knockout-first, or gene-trap knockout allele. Interestingly, many tm1a gtKO mouse models may still express residual wild-type mRNA and protein, producing viable homozygous mice that otherwise would not be obtained via a classic knockout (null) approach. Indeed, using this gtKO strategy, we obtained viable homozygous *Scn2a*^*gtKO/gtKO*^ mice that survive to adulthood. We confirm a significant reduction in *Scn2a* expression and examine the health, sensory, and innate behavior of gtKO *Scn2a* mice. We show that *Scn2a*^*gtKO/gtKO*^ mice may be a valid mouse model for in-depth studies of disorders associated with *SCN2A* deficiency.

**Figure 1.**
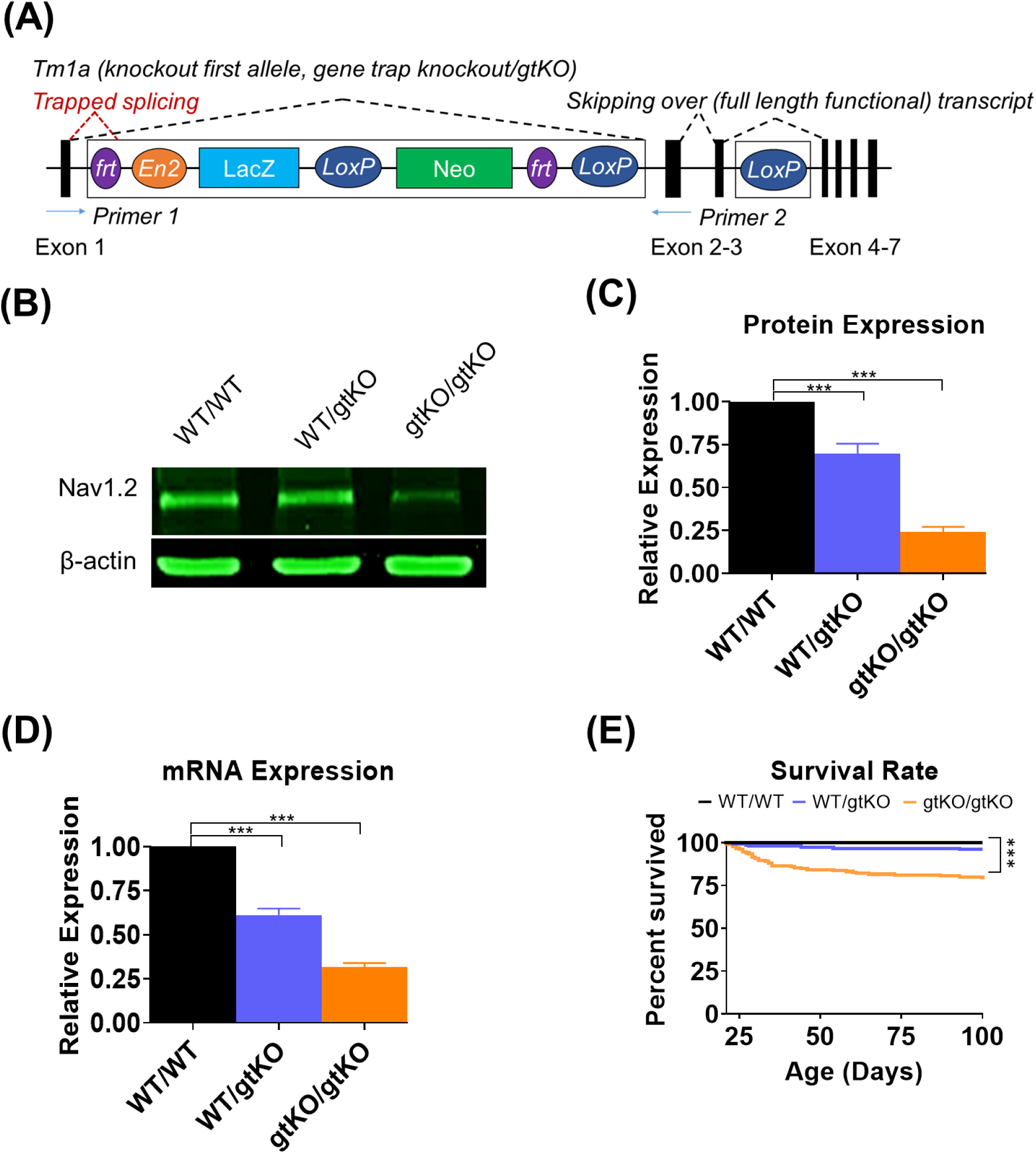
Substantial reduction of Nav1.2 expression in Scn2a^gtKO/gtKO^ mice. **A)** Diagram of the tm1a gene trap cassette between Exon 1 and 2. Abbreviations: Frt, Flp recognition target (purple); En2, engrailed-2 splice acceptor (orange); LacZ, lacZ β-galactosidase (light blue); LoxP, locus of X-over P1 (dark blue); and Neo, neomycin (green). **B)** Representative Western Blot with β-actin control. **C)** Reduced expression levels of Nav1.2 in heterozygous and homozygous gene trap knock-out (gtKO) mice normalized to WT littermates. **D)** Expression levels of *Scn2a* are reduced in both heterozygous and homozygous gtKO mice normalized to WT littermates. **E)** Survival rate to 100 days old is reduced in *Scn2a*^*gtKO/gtKO*^ mice but still high.

## MATERIAL AND METHODS

### Animals

C57BL/6N-*Scn2a1*^*tm1aNarl*^/Narl (here we referred as *Scn2a*^*WT/gtKO*^) mice were generated from the National Laboratory Animal Center Rodent Model Resource Center (Taipei City, Taiwan) based on a modified gene-trap design [14]. All animal experiments were approved conducted in accordance with the Institutional Animal Care and Use Committee (IACUC) and the National Institutes of Health Guide for the Care and Use of Laboratory Animals. Heterozygous (HET, *Scn2a*^*WT/gtKO*^) mice were used as breeding pairs. Experiments were performed on mice that were at least six generations from the founders. WT C57BL/6N were used to back-cross every few generations. Pups were weaned 21-28 days after birth and given a nutritionally fortified dietary gel supplement (DietGel 76A; ClearH_2_O, Portland, ME) to reduce diarrhea and improve hydration. Mice were same-sex housed in mixed-genotype groups (3-5 mice per cage) on vented cage racks with 1/8” Bed-o-cobb bedding (Anderson, Maumee, OH, USA) and >8g of nesting material as enrichment (shredded paper, crinkle-cut paper, and/or cotton nestlet) on a 12hr light cycle. Food (2018S Teklad from Envigo) and reverse osmosis water was given ad lib.

Mice (3-5 months old) were habituated to the behavior room 45-60 minutes before testing. No more than two behavioral tests were performed each week on the same mouse. A minimum of 6 of each sex of each genotype were tested. All work was performed at the Purdue Behavioral Core during the dark cycle.

### Genotyping and molecular analysis

Mice were genotyped at wean (21-28 days) via ear punch. Genotyping for the tm1a cassette was performed using gene-specific polymerase chain reaction (PCR) on DNA extracted from ear tissue using a tissue DNA extraction kit (Macherey-Nagel, Bethlehem, PA, USA) with primers suggested by the stock facility (forward: GAGGCAAAGAATCTGTACTGTGGGG, reverse: GACGCCTGTGAATAAAACCAAGGAA). PCR product for the wild type cassette is 240 base pairs (bp) and the tm1a allele is 340 bp. qPCR was performed on whole brains using an RNA extraction kit (Qiagen, Hilden, Germany) and a cDNA kit (ThermoFisher). Primers for qPCR were ATTTTCGGCTCATTCTTCACACT (forward) and GGGCGAGGTATCGGTTTTTGT (reverse). Western Blot was performed using an anti-Scn2a antibody (Alomone, Jerusalem, Israel) and a β-actin control (Cell Signaling Technology, Danvers, MA, USA).

### Metabolism

Mice were singly housed in a metabolism monitoring system (Techniplast, West Chester, PA, USA) in their home room. System consists of a water bottle holder, side food hopper, and a wire floor to allow fecal matter and crumbs to fall through while urine funnels to a different collection tube. Fecal boli was separated from the food crumbs. 20g of standard food was provided with water ad lib. Plastic huts were used in lieu of nesting material. Measurements (weight of food and crumbs, number and weight of fecal boli) were taken daily and averaged for each animal as its final value.

### SHIRPA

A modified SmithKline Beecham/Harwell/Imperial College/Royal London Hospital Phenotype Assessment (SHIRPA) [16] was conducted as a preliminary phenotypic screen. Genotype-blind researchers give a score (0-5) to home cage and transferring observations using criteria (**Table 1**). Body position, spontaneous activity, and tail posture were assessed after 5 minutes of home cage observation. Then mice were removed from the cage by holding by the base of the tail and trunk curl, limb grasping, and postural reflex was assessed. Next mice were placed so that their front paws were gripping the end of the wire food hopper to measure grip strength. When mice were slowly lowered back in their cage, extension was observed. On a different day, mice were removed and replaced on their back to assess righting reflex.

**Table 1.** SHIRPA Criteria.

| Name | 0 | 1 | 2 | 3 | 4 | 5 |
| --- | --- | --- | --- | --- | --- | --- |
| Body position | Completely flat (belly on ground, muscles tensing as to flatten out as much as possible) | Lying prone (belly on ground but no muscle tensing) | Ball (mouse is spherical shaped, huddled) | Hunched (body extended but back and neck are bent) | Sitting or standing | Rearing (on back legs) |
| Spontaneous Activity | None, resting (no movement for >5 min) | Casual scratch and groom, slow movement (movement for <1 min) | Vigorous scratch and groom, moderate movement (movement for >2 min) | Vigorous, rapid/dart movement (movement for >3 min) | Extremely vigorous, rapid/dart movement (constant movement for >2 min, movement for >3 min total) | Rapid movement and jumping (constant movement for >4 min, movement for >4 min total) |
| Tail Posture | No attempt to control | Movement attempted, but unsuccessful | 45 degrees down | Horizontally extended, parallel with the floor | Elevated >90 degrees | Active, seemingly random movement |
| Trunk Curl | None | ¼ curl | ½ curl | ¾ curl | Full curl | Hyperextended curl |
| Limb Grasping | None | Some attempt by <2 limb for <3 seconds | Flailing by all limbs for <5 seconds | Flailing by <2 limbs for 5-10 seconds | Flailing by all limbs for 5-10 seconds | Toes extended and feet flailing continuously |
| Extension (when lowering) | None | Upon nose contact | Upon whisker contact | Before whisker contact (18mm) | Early extension (25mm) | Anticipated extension (above cage) |
| Grip | None | Slight grip attempted, not effective | Slight (<2 paws) grip, semi-effective | Moderate (>2 paws) grip, effective | Active grip (2-3 paws), effective | All paws, effective |
| Postural reflex | No change in posture | Clasping of hind and forepaws, head straight down or by paws | Clasping of hind paws and forepaws, head raise level with ground | Curling inward of paws and or/head raised level with ground | Spreading of hind limbs | Spreading of hind limbs and head raised to be level with the ground |
| Righting reflex | No attempt | Failed attempt | All feet back on the ground unassisted within 5 seconds | N/A | N/A | N/A |

### Olfactory sensitivity and male sexual preference

Olfactory sensitivity was tested as described previously [17]. Mice were placed in a standard cage bottom with no bedding for 5 minutes. After habituation, filter paper covered in 1% cinnamon extract (McCormick & Company, Sparks Glencoe, MD, USA) at one end and water at the other. For male sexual preference test, sexually naïve males were habituated as above and tested for partner preference. Protocol was modified from a similar test [18]. Filter paper containing urine (diluted 1:5 in RO water) from a male breeder and an estrus female (estrus confirmed through visual observation and microscope slide analysis) were placed at either end.

### Quinine Avoidance

The quinine avoidance test was conducted using a previously defined protocol [19]. Mice were habituated to two 50mL conical bottles with sippers on them for 10 days (both water) in a two-grommet cage. After habituation, the bottles were replaced with one 0.6mM quinine in reverse osmosis (RO) water and one of just RO water on random sides alternating daily for four days. The weight was recorded daily at the same time each day. A cage without a mouse was used as a control to measure daily drip rate, which was subtracted from the total volume to obtain the total volume drank.

### Auditory startle

Auditory startle was conducted in SR-LAB startle response chambers (San Diego Instruments, San Diego, CA, USA) based on previous studies [20] [21] [22]. Mice were habituated to the container and 70dB background white noise for 5 min. Then eight blocks of five 40msec stimuli (75, 85, 95, 105, 115 dB) at 4kHz were presented in a pseudo-random order with a random inter-stimulus interval between 10-30 sec (mean 20 sec) with continuous background white noise of 70 dB throughout the experiment. Total session time was approximately 15 min. Data was analyzed using a two-way repeated measures ANOVA with genotype and volume.

### Thermal Response

For heat tolerance, mice were placed on a 50°C hot/cold plate system model PE34 (IITC Life Sciences, Woodland Hills, CA, USA). Cold tolerance was at 7.5°C. Dynamic hot plate thermal sensitivity started at 30°C and increased to 60°C at a rate of 2°C/sec. Dynamic cold plate cold sensitivity started at 30°C and decreased to 5°C at a rate of 2°C/sec. Temperatures were determined based on previous studies [23] [24] [25]. Time (sec) until distress was recorded using a stopwatch of a genotype blind observer. Distress was defined as flicking, biting, or licking of the arms, legs, or tail. Animals were removed at first sign of distress or after 20 seconds if no signs were observed.

### von Frey Mechanical Sensitivity

A simplified up-down von Frey test was performed as previously described [26]. Mice were placed in a sectioned chamber with a mesh bottom for 5 minutes to habituate. von Frey fibers (BioSeb Lab Instruments, Vitrolles Cedex, France), starting with 2.0g, were pressed on the mouse’s hind paw pad until fiber started to bend and held for three seconds. A response would be a withdrawal, flicking, or flinching. If there was a response, a thinner fiber was applied. If no response, then a thicker fiber was applied. This was repeated three times with not exceeding 4.0g.

### Nesting Behavior

Nesting behavior was assessed using a modified protocol [27]. Mice were singly housed in clean cages two hours before start of dark cycle with bedding and *ad lib* food and water. One 5cm by 5cm square condensed cotton nestlet (Ancare, Bellmore, NY, USA) was added to the front right corner of the cage one hour before start of dark cycle. Height and quality of the nest were recorded at 36 hours after addition of the nestlet. Height was recorded as the average of the four corners of the nest. Nest quality score was the average score rated by three genotype-blind researchers based on a 0-5 point scale according to the following criteria: **0** = nesting material has not been touched; **1**= nesting material is largely untouched (>90% intact); **2** = nesting material is partially torn up (50-90% remaining intact); **3** = nesting material is mostly shredded but often there is no identifiable nest site: < 50% of the nesting material remains intact but < 90% is within a quarter of the cage floor area, i.e. the cotton is not gathered into a nest but spread around the cage (the material may sometimes be in a broadly defined nest area but the critical definition is that 50-90% has been shredded); **4** = an identifiable, but flat nest: > 90% of the nesting material is torn up, the material is gathered into a nest within a quarter of the cage floor area, but the nest is flat, with walls higher than mouse body height (curled up on its side) on less than 50% of its circumference; **5** = a (near) perfect nest: > 90% of the nesting material is torn up, the nest is a crater or cocoon shaped with defined walls higher than 50% of the mouse’s circumference.

### Marble Burying

Marble burying test was modified from previous protocols [28]. 15 black glass marbles arranged 3×5 were placed in a clean home cage with 5cm deep bedding. Number of marbles at least two-third buried in a 30-minute session was recorded by three blind investigators then averaged for that mouse’s score.

### Statistical analysis

Data was analyzed using a two-way analysis of variance (ANOVA) to identify any genotype, sex, or genotype x sex interaction effects then analyzed using a post-hoc with Bonferroni corrections unless noted otherwise. Olfactory sexual preference was analyzed using a one-way ANOVA since only males were studied. Weight was also analyzed using one-way ANOVA since there is an inherent sex difference. Computer software GraphPad Prism 8.3.0 (GraphPad Software Inc, La Jolla, CA, USA) was used for statistical analysis and figure generation. Results are presented as mean ± S.E.M. Significance was set at p≤0.05 (*), p≤0.01 (**), and p≤0.001 (***).

## RESULTS

### Validation of gene-trap knockout of Scn2a

Here we characterize a *Scn2a* partial loss-of-function mouse model designed with a modified targeted gene trap (gtKO) strategy [14]. gtKO of *Scn2a* is achieved by inserting a gene trap into Intron 1 of the *Scn2a* gene, thus “trapping” the splicing of the first exon of *Scn2a* exons to produce a LacZ fusion transcript and truncating the wild-type transcript (**Figure 1A**). In theory, since no exon sequences are removed, residual wild-type mRNA and protein could be expressed via splicing around the trapping cassette. However, the extent of wild type transcript and protein may vary and must be determined empirically.

Using two primers in exons flanking the inserted trapping cassette, which capture the “skip over” full length transcripts, we sequenced RT-PCR products and indeed identified transcripts containing both full-length Exon 1 and Exon 2 sequences supporting the notion that full length non-truncated *Scn2a* transcripts exist. We also identified transcripts containing Exon 1 spliced to LacZ sequences of the trapping cassette, confirming that gene trapping is functioning, but does not completely eliminate wild-type transcripts.

To measure the residual amount of *Scn2a*, we assayed both mRNA and protein expression of *Scn2a* in gtKO mice. Compared to wild type (WT, *Scn2a*^*WT/WT*^) expression in whole brain, *Scn2a* protein levels in heterozygous *Scn2a*^*WT*/gtKO^ brain is 69.6±5.9% and 24.02±3.1% in *Scn2a*^*gtKO/gtKO*^ brain compared to WT which had a significant genotype main effect (F_2,12_=80.31, p<0.0001), but not any sex (F_1,12_=0.1111, p=0.7447) or genotype x sex effects (F_2,12_=0.0829, p=0.0920) (**Figure 1B and 1C**). Similarly, the mRNA level in heterozygous (HET, *Scn2a*^*WT/gtKO*^) mice is 61.2±3.6% whereas the mRNA level in gtKO homozygous (HOM, *Scn2a*^*gtKO/gtKO*^) is only 31.8±2.2% which is a significant genotype effect (F_2,6_=112.3, p<0.001) with no significant sex (F_1,6_=0.4287, p=0.5369) or interaction effects (F_2,6_=1.244, p=0.3531) (**Figure 1D**). Thus our molecular data supports the notion that the gtKO allele is not null, but rather expresses wild-type *Scn2a* protein at about a quarter of the normal level of expression.

### Maintenance behaviors

Heterozygous mice were used as mating pairs. According to Mendelian genetics, the theoretical ratio of a *Scn2a*^*WT/gtKO*^ x *Scn2a*^*WT/gtKO*^ breeding pair is 1:2:1 (WT: HET: HOM). Our mice colony of 839 pups across 100 litters had an average rate of 1:1.65:0.89 (WT: HET: HOM) with no sex differences. A soft, high-calorie diet was provided at wean to reduce mortality of *Scn2a*^*gtKO/gtKO*^ due to failure to thrive. The survival rate to 100 days was 100% for WT, 96.0% for *Scn2a*^*WT/gtKO*^, and 78.4% for *Scn2a*^*gtKO/gtKO*^ (**Figure 1E**). *Scn2a*^*gtKO/gtKO*^ mice died (or were euthanized per IACUC guidelines) as a result of severe diarrhea, penile prolapse, rectal prolapse, or hydrocephalus. At 100 days, the body weights were significantly different by genotype (F_2,53_=37.83, p<0.0001), sex (F_1,53_=113.7, p<0.0001), and the interaction of genotype x sex (F_2,53_=7.591, p=0.0013). Interestingly, female *Scn2a*^*WT/gtKO*^ and *Scn2a*^*gtKO/gtKO*^ weighed significantly less than WT (post-hoc p=0.0014 HET, p<0.0001 HOM). On the other hand, only male *Scn2a*^*gtKO/gtKO*^ weighed significantly less (p<0.0001) whereas male *Scn2a*^*WT/gtKO*^ weighed similar to WT (p>0.9999) (**Figure 2**). The decreased survivability and reduced weight of *Scn2a*^*gtKO/gtKO*^ mice, possibly due to gastro-intestinal complications and failure to thrive, steered us to characterize their basic metabolism.

**Figure 2.**
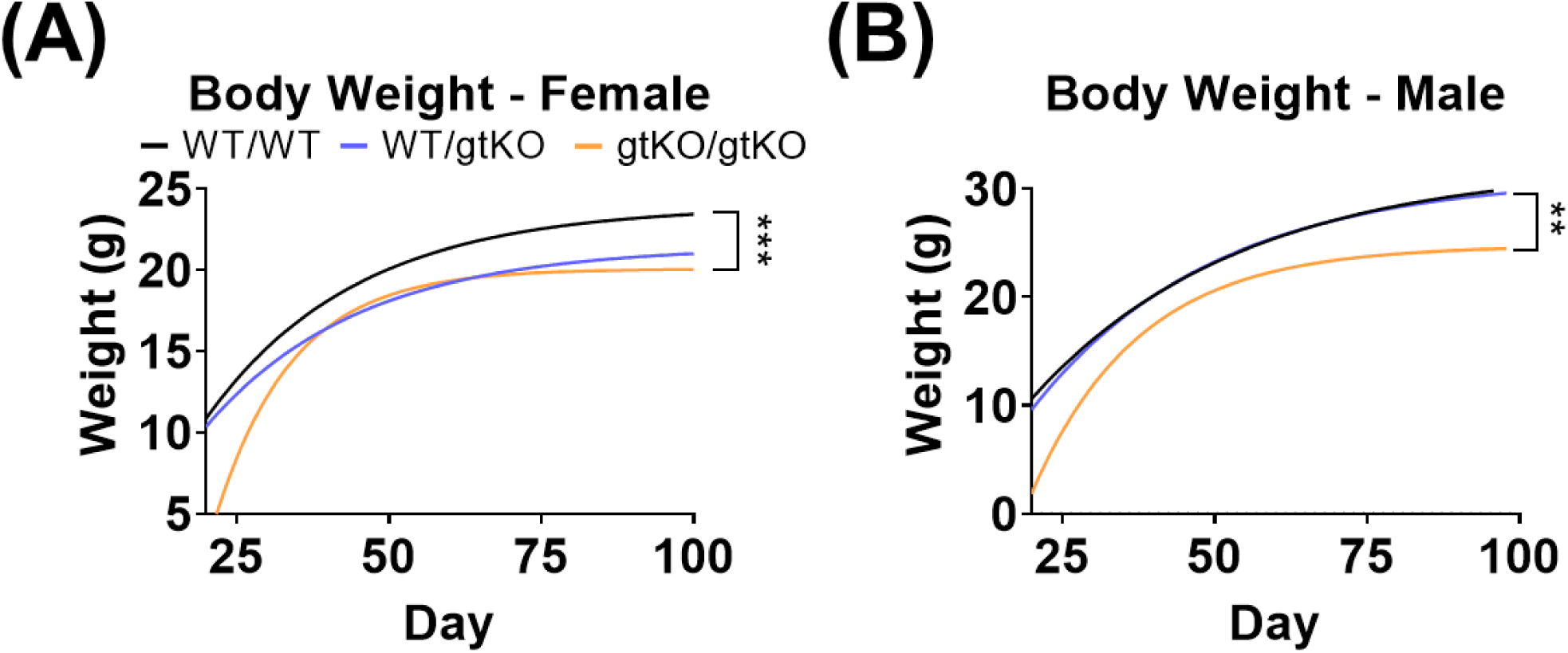
Scn2a^gtKO/gtKO^ mice have significantly smaller body weights. **A)** Female body weight with a one phase decay least squares line of best fit with rate constants of 0.0405, 0.0396, and 0.07892 with R^2^ of 0.7971, 0.7432, and 0.8815 for WT, *Scn2a*^*WT/gtKO*^, and *Scn2a*^*gtKO/gtKO*^ respectively. **B)** Male body weight with a one phase decay least squares line of best fit with rate constants of 0.0290, 0.0333, and 0.0569 with R^2^ of 0.9181, 0.9032, and 0.9228 for WT, *Scn2a*^*WT/gtKO*^, and *Scn2a*^*gtKO/gtKO*^ respectively.

During temporary opposite-sex breeding husbandry, we observed that homozygous mice did not mate despite trying multiple paradigms of breeding pairs: female *Scn2a*^*gtKO/gtKO*^ x male *Scn2a*^*gtKO/gtKO*^, female *Scn2a*^*gtKO/gtKO*^ x male WT, and male *Scn2a*^*gtKO/gtKO*^ x female WT. Specifically, male *Scn2a*^*gtKO/gtKO*^ had no mounting behavior with female WT in estrus. Since we observed that the *Scn2a*^*gtKO/gtKO*^ mice have little to no mating activity, we conducted a socio-sexual olfactory investigatory preference test of male mice between urine from an estrus female and urine from a male breeder. The *Scn2a*^*gtKO/gtKO*^ mice did not have a preference for the female scent over the male scent (post-hoc p=0.2048) but *Scn2a*^*WT/gtKO*^ showed a preference for the female over the male similar to WT (p>0.9999) (**Figure 3A**).

**Figure 3.**
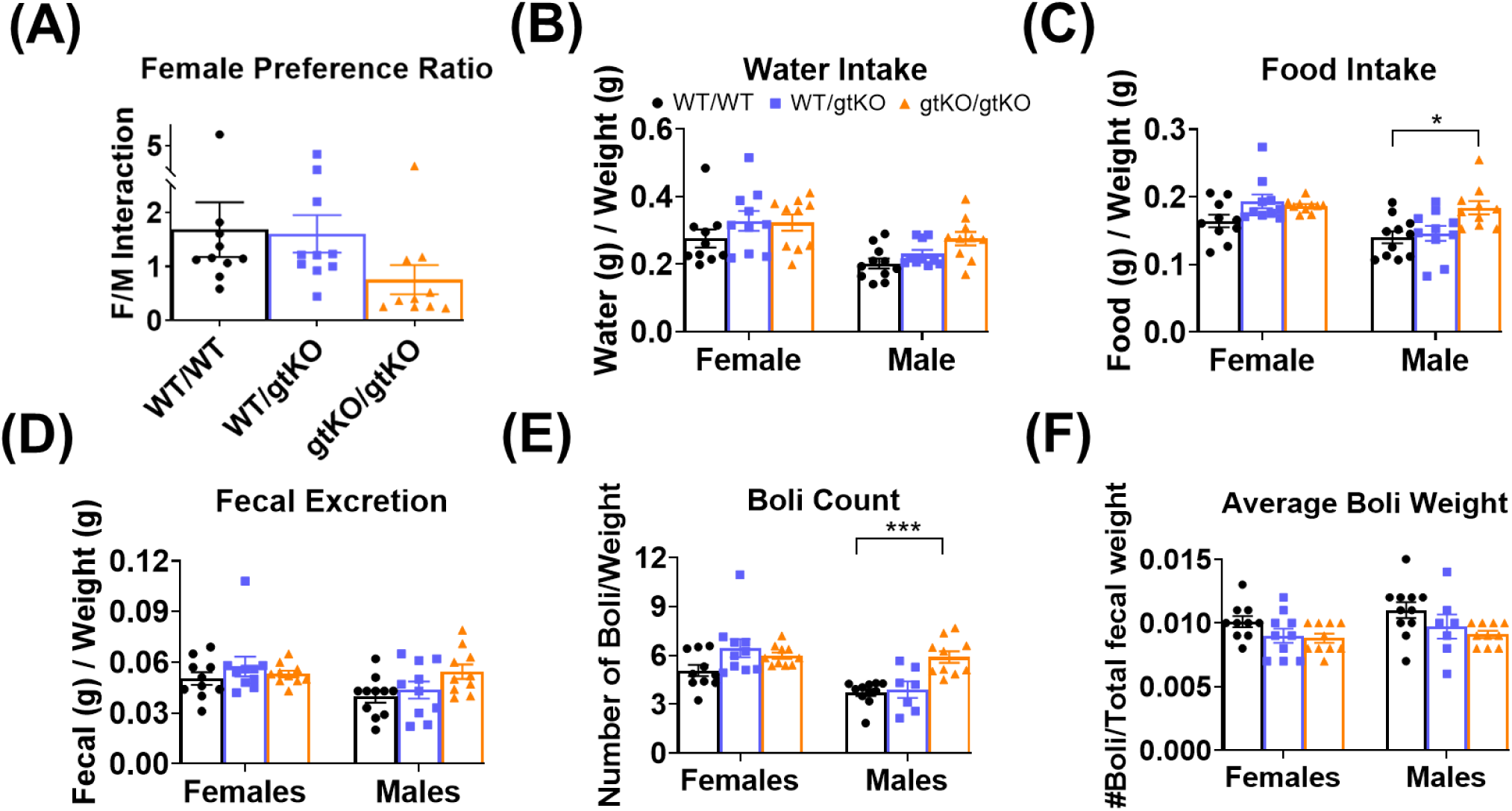
Scn2a^gtKO/gtKO^ mice have impairments in mating and metabolism. **A)** Quantification of preference for estrus female over male urine interaction ratio. Female/male (F/M) ratio of 1 means no preference, >1 means preference for the female stimulus, and <1 means preference for the male stimulus. **B)** Water intake is slighlty increased in *Scn2a*^*gtKO/gtKO*^ mice. **C)** Food intake is significantly increased in male *Scn2a*^*gtKO/gtKO*^ mice. **D)** Male *Scn2a*^*gtKO/gtKO*^ mice have a slight increase fecal excretion. **E)** Male *Scn2a*^*gtKO/gtKO*^ mice also have a significant increased fecal boli count. **F)** *Scn2a*^*WT/gtKO*^ and *Scn2a*^*gtKO/gtKO*^ mice of both sexes have a slight decreased boli weight.

We monitored the animal’s food and water intake and compared that to its fecal production. Data was normalized based on the individual’s weight. The water intake varied with genotype (F_2,54_=4.015, p=0.0236) and sex (F_1,54_=16.79, p=0.0001) but not sex x genotype interaction (F_2,54_=0.6507, p=0.5256) (**Figure 3B**). Food intake significantly increased with genotype (F_2,54_=6.603, p=0.0027) with a significant sex effect (F_2,54_=11.02, p=0.0016) but no significant interaction effect (F_2,54_=2.967, p=0.0597) (**Figure 3C**). We did not observe a significant genotype effect in fecal excretion (F_2,54_=2.195, p=0.1210) (**Figure 3D**) but boli count had a significant genotype (F_2,54_=8.994, p=0.0004), sex (F_1,54_=8.994, p=0.0004), and interaction effect (**Figure 3E**). The average boli weight significantly varied by genotype (F_2,54_=5.131, p=0.0093) (**Figure 3F**). Specifically, post-hoc analysis revealed that male *Scn2a*^*gtKO/gtKO*^ mice had a significantly increased food intake compared to male WT (p=0.0163). Consequently, male *Scn2a*^*gtKO/gtKO*^ mice had a significantly increased fecal boli count compared to male WT (p=0.0012). Together, these results suggest that male *Scn2a*^*gtKO/gtKO*^ mice have significant differences in eating and waste excretion compared to WT.

For general behavior, a preliminary SmithKline Beecham/Harwell/Imperial College/Royal London Hospital Phenotype Assessment (SHIRPA) was conducted. The SHIRPA is a qualitative analysis (rated on a scale of 0-5, or 0-3 for righting reflex) of basic behavior that can provide direction for follow-up experiments. Of the nine behaviors observed, body position, trunk curl, grip, and postural reflex were significantly different between WT and *Scn2a*^*gtKO/gtKO*^ mice after post-hoc analysis. Specifically, *Scn2a*^*gtKO/gtKO*^ mice had a lower body position rating (more hunched), less trunk curl when held by the tail, decreased grip ability (slight grip attempted, not effective), and less of a postural reflex when held by the tail (some head raise, front paws curled inward, some spread of hind limbs). Activity, tail posture, limb grasping, extension, and righting reflex were similar between WT, *Scn2a*^*WT/gtKO*^, and *Scn2a*^*gtKO/gtKO*^ mice (**Supplemental File 1**). There were no significant sex differences in any of the nine SHIRPA parameters.

### Locomotion and sensory abilities

We used open field locomotion to assess for motor deficits. Total distance traveled had a significant genotype (F_2,39_=3.517, p=0.0394), sex (F_1,39_=5.568, p=0.0234), and interaction effect (F_2,39_=5.244, p=0.0096). Post-hoc analysis revealed that the total distance traveled was significantly increased for female *Scn2a*^*gtKO/gtKO*^ (p=0.0103) but not female *Scn2a*^*WT/gtKO*^ (p>0.9999) when compared to female WT. There was no genotype difference between the males (F_2,18_=0.4349, p=0.6539). Based on this general motor test, we suggest that female *Scn2a*^*gtKO/gtKO*^ mice display hyperactivity.

We further tested the sensory modalities of this mouse model. Olfactory sensitivity was assessed by comparing the time spent interacting (i.e. sniffing, licking) with a cinnamon solution versus water as described previously [17] as a way to test olfactory functionality. There was no significant genotype difference in cinnamon preference (F_2,30_=1.824, p=0.1788) (**Figure 4A**) or total interaction time (F_2,30_=0.3776, p=0.6887) (**Figure 4B**). These olfactory results show that *Scn2a*^*gtKO/gtKO*^ mice have a normal smelling ability similar to WT. Quinine avoidance was used as a taste test paradigm in which the mice had two bottles, one with their normal reverse osmosis water and the other with quinine (bitter taste). There were no genotype differences in quinine avoidance (F_2,30_=2.463, p=0.1022), suggesting that these *Scn2a*^*gtKO/gtKO*^ mice do not have a deficit in discriminating bitter tastes (**Figure 4C**). Hearing was tested using acoustic startle at five volumes ranging from 75-115dB at the same frequency (4kHz). There was a significant difference in genotype at 75dB (F_2,30_=4.972, p=0.0136) but no other significant genotype effects at the other volumes (**Figure 4D**).

**Figure 4.**
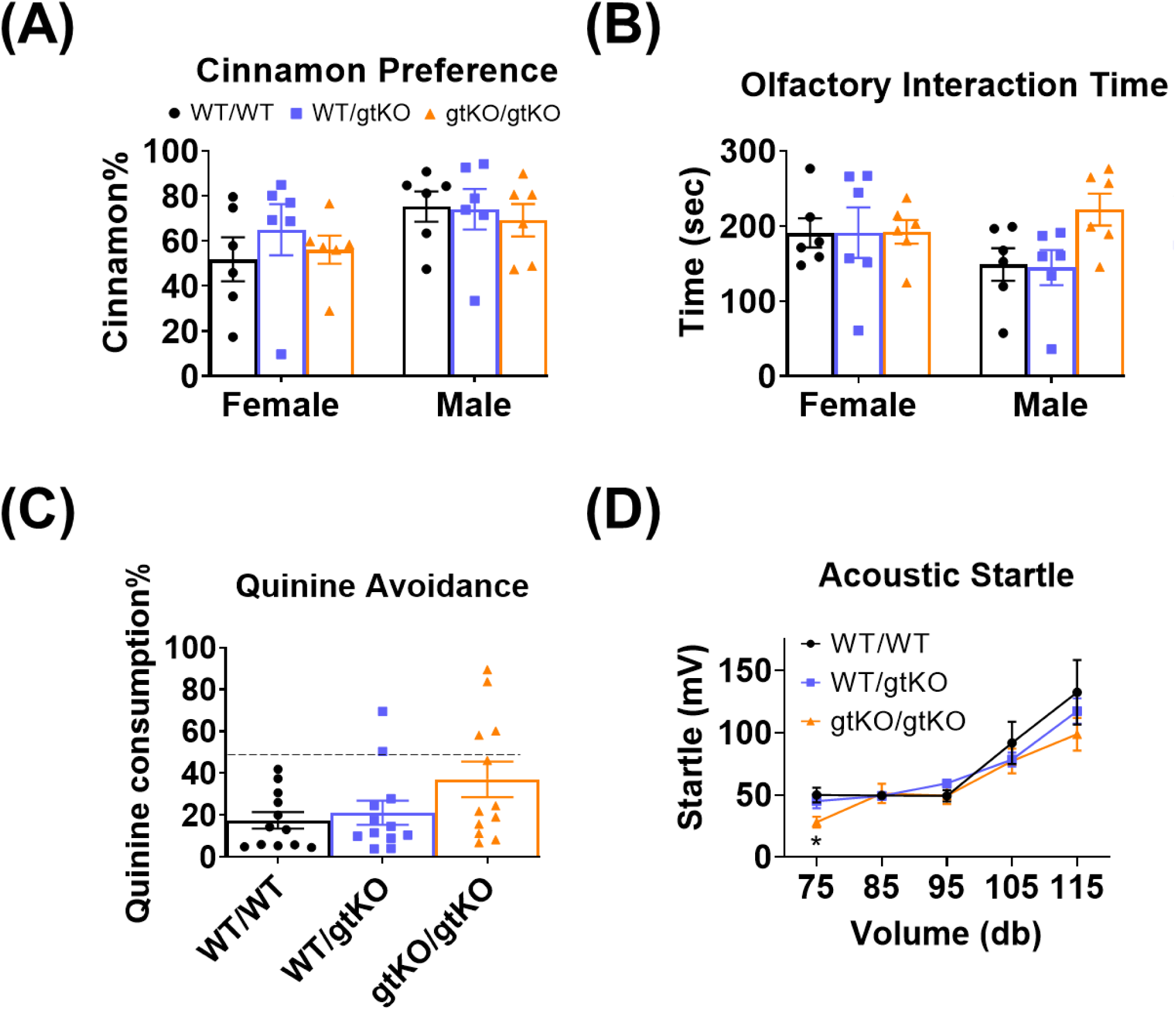
Scn2a^gtKO/gtKO^ mice do not have any major effects on sensory systems. **A)** All mice have a similar preference for cinnamon over water. **B)** Male *Scn2a*^*gtKO/gtKO*^ mice have a slight increase in interaction time with both cinnamon and water stimuli. **C)** *Scn2a*^*gtKO/gtKO*^ mice have a slight decrease in quinine avoidance (increased quinine consumption). **D)** There is no genotype difference in acoustic startle overall, but a significant decrease in *Scn2a*^*gtKO/gtKO*^ mice startle at 75dB.

To test the response to thermal and cold stimuli, mice were placed on temperature-controlled plates. We tested both temperature tolerance and sensitivity. We define tolerance as the latency of response on a plate of static temperature, whereas sensitivity is the latency of response to scaled temperatures. The thermal/cold tolerance test was a plate kept at a static temperature (50°C for hot and 7.5°C for cold). Genotype differences were significant in both the hot (F_2,30_=6.102, p=0.0060) and the cold (F_2,30_=31.29, p<0.0001) static plate paradigms. Specifically, post-hoc analysis found that *Scn2a*^*gtKO/gtKO*^ (p<0.0001) and *Scn2a*^*WT/gtKO*^ (p=0.0098) had a significant decreased tolerance for cold; whereas only *Scn2a*^*gtKO/gtKO*^ (p=0.0082) and not *Scn2a*^*WT/gtKO*^ (p>0.9999) had a significant decreased tolerance for heat (**Figure 5A and 5B**). Thermal/cold sensitivity was a dynamic hot/cold plate that started at 30°C and ramped (2°C/sec) to the desired temperature (60°C for hot and 5°C for cold). No significant genotype effects were detected in hot (F_2,30_=0.3096, p=0.7360) or cold sensitivity (F_2,30_=2.015, p=0.1510) (**Figure 5C and 5D**). Mechanical sensitivity was measured by applying von Frey fibers to the hindpaw pad using an up-down protocol until the mouse responded to the fiber. There were no mechanical sensitivity differences between genotypes (F_2,54_=0.0249, p=0.9755) or sex (F_1,54_=0.0750, p=7853). Therefore, we conclude that *Scn2a*^*gtKO/gtKO*^ mice have a significant decrease in thermal and cold tolerance, but not thermal, cold, or mechanical sensitivity.

**Figure 5.**
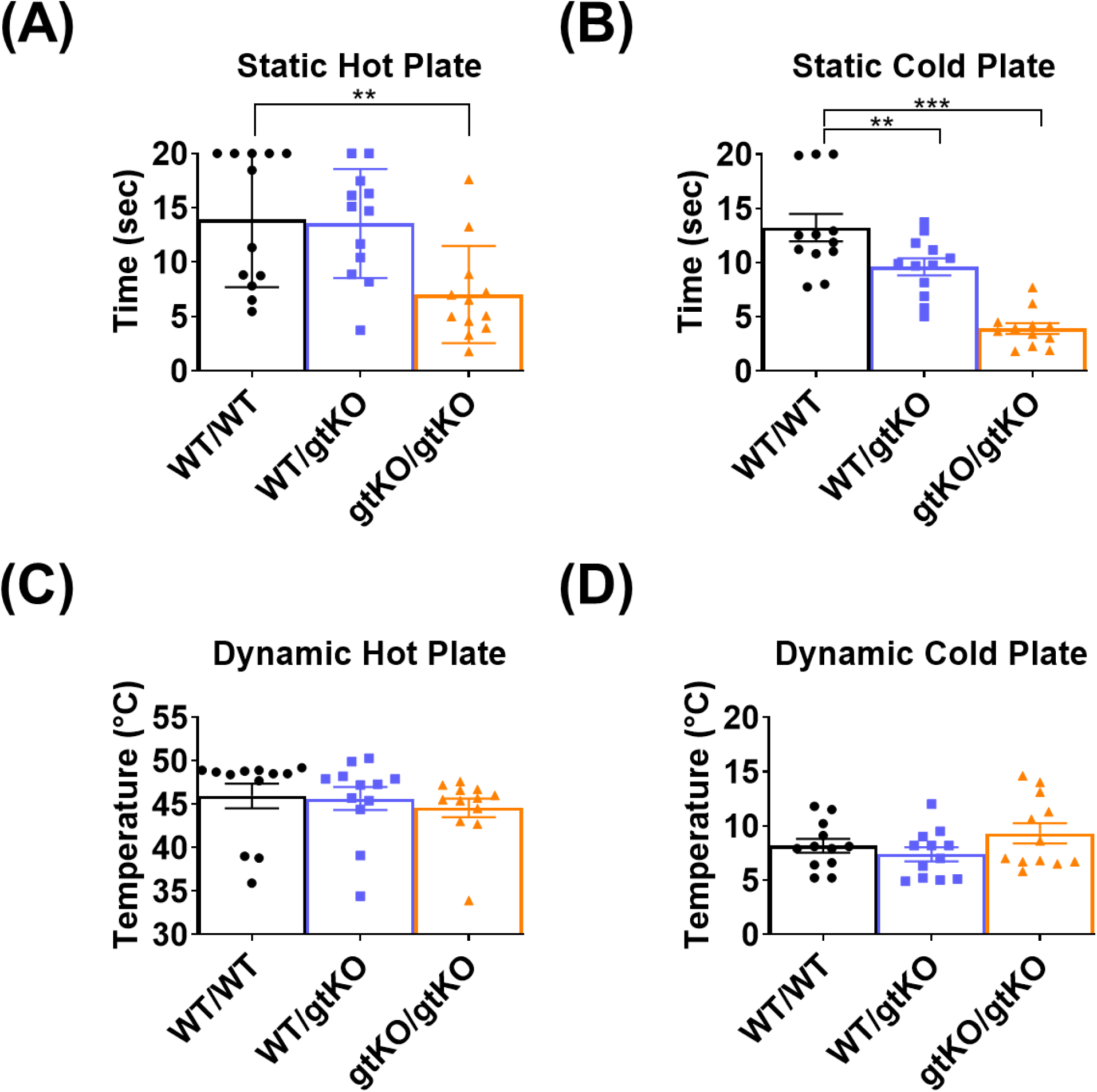
Scn2a^gtKO/gtKO^ mice have deficits in thermal tolerance but not thermal or mechanical sensitivity. **A)** Response to a static hot plate was significantly faster in *Scn2a*^*gtKO/gtKO*^. **B)** Both *Scn2a*^*WT/gtKO*^ mice and *Scn2a*^*gtKO/gtKO*^ mice had a decreased latency to respond to the cold plate. **C)** Scaled temperature from 30°C to 55°C on the hot plate showed similarity among all genotypes. **D)** *Scn2a*^*gtKO/gtKO*^ mice had a slight increase in cold sensitivity during the scaled dynamic cold plate of decreasing temperature from 20°C to 0°C.

### Innate behaviors

Nesting is a thermoregulatory and social innate behavior of mice [27]. Significant genotype effects in nest quality (F_2,39_=155, p<0.0001) (**Figure 6B**) and height (F_2,39_=36.12, p<0.0001) (**Figure 6C**) were observed. Importantly, *Scn2a*^*gtKO/gtKO*^ mice have little to no nesting activity, having significantly less nest quality (p<0.0001 males and females) as rated on a scale of 0-5 (see **Materials and Methods**) and a much smaller nest height (p=0.0004 females, p<0.0001 males) from post-hoc analysis separated by sex. On the other hand, *Scn2a*^*WT/gtKO*^ mice had a nest similar to WT in both quality (p>0.9999 males and females) and height (p>0.9999 males and females). Marble burying can also be considered as innate behavior [29]. In the marble burying test, a significant genotype (F_2,30_=130.90, p<0.0001) and sex difference (F_1,30_=6.6720, p=0.0149) were observed. *Scn2a*^*gtKO/gtKO*^ mice buried little to no marbles (p<0.0001 for males and females), while *Scn2a*^*WT/gtKO*^ buried a similar number to WT (p>0.9999 for males and females) (**Figure 7**). Thus we concluded that *Scn2a*^*gtKO/gtKO*^ mice have abnormal innate behavior of nest building and marble burying.

**Figure 6.**
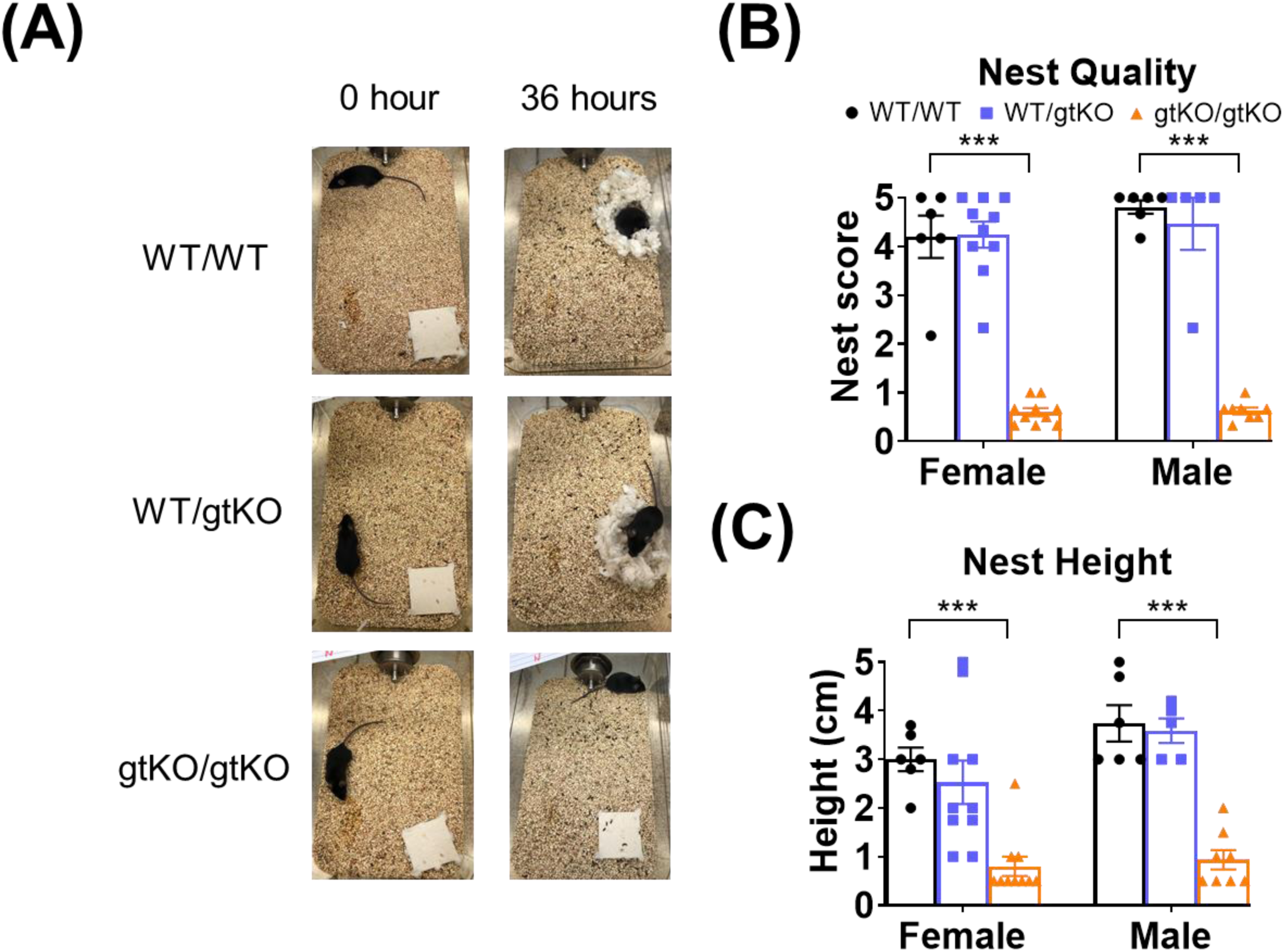
Scn2a^gtKO/gtKO^ mice have little to no nesting behavior or male sexual preference. **A)** Representative nests after 36 hours. Top right image (WT after 36 hours) shows an example of a quality rating of 5 (on a scale from 0-5) or near perfect nest with all of the nesting material in a crater-shape with walls higher than the mouse. Bottom right image (*Scn2a*^*gtKO/gtKO*^ mice after 36 hours) shows a quality rating of 0 with little to no nesting material disturbed. **B)** Quantification of nest quality reveals both sexes of *Scn2a*^*gtKO/gtKO*^ mice have significantly decreased nesting quality. **C)** Nest height is also significantly decreased in both sexes of *Scn2a*^*gtKO/gtKO*^ mice.

**Figure 7.**
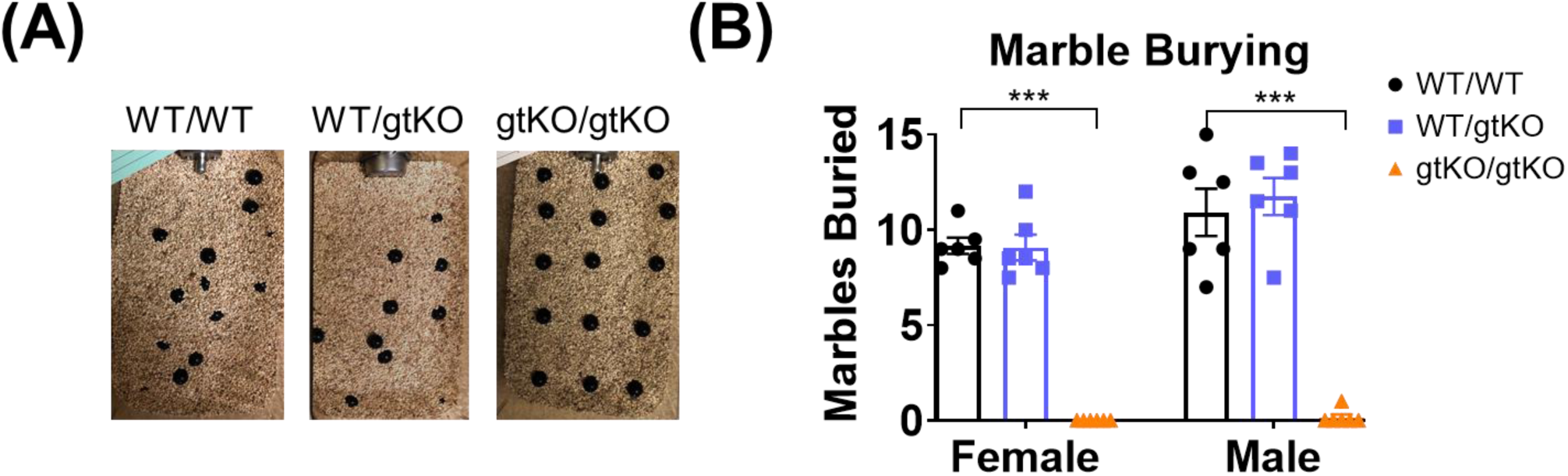
Scn2a^gtKO/gtKO^ mice have little to no marble burying behavior. **A)** Representative marble buried. 15 marbles are arranged in a 3×5 grid in their standard home cage with 5cm of bedding. Marbles were counted after 30 minutes and counted buried if >2/3rds was under the bedding. **B)** Both male and female *Scn2a*^*gtKO/gtKO*^ bury significantly less marbles than *Scn2a*^*WT/gtKO*^ mice and WT mice.

## DISCUSSION

In this paper we characterized a hypomorphic allele of *Scn2a* in mice (*Scn2a*^*gtKO*^) and provide evidence to suggest that *Scn2a*^*gtKO/gtKO*^ mice could serve as model for *SCN2A* loss-of-function channelopathies. Our study shows that *Scn2a*^*gtKO/gtKO*^ mice have little-to-no mating, nesting, or marble burying activity, indicating decreased innate behavior. *Scn2a*^*gtKO/gtKO*^ mice have normal olfactory, taste, hearing, and mechanical sensitivity but a decrease in thermal and cold tolerance. Our *Scn2a*^*gtKO*^ model can be used to further understand the etiology of *SCN2A*-related disorders in humans and to develop and test treatments.

*Scn2a* encodes for voltage-gated sodium channel Nav1.2, which is mainly responsible for action potential firing in the central nervous system throughout life. Structure-function studies of Nav1.2 have identified genetic “hotspots” that have an increased number of disorder-associated variants in the voltage sensing domain, intracellular terminal domains, and pore loops of the ion selectivity filter [7]. The genotype-phenotype of *Scn2a* dysfunction in humans is highly correlative because patients with the same genetic variants present similar phenotypic features [30] [31]. The severity of clinical phenotypes also correlates with the degree of neuronal electrophysiological dysfunction [32]. Our gene trap knockout *Scn2a* mouse model (*Scn2a*^*gtKO/gtKO*^ mice) serves as a model to understand *SCN2A* deficiency, which could result from a nonsense mutation, loss-of-function mutation or splice deficit.

Mice models have been a cornerstone for our understanding of *in vivo* biological mechanisms and have provided valuable insights for many genetic disorders. Mechanistic investigations to gain critical knowledge of *SCN2A* function at the cellular, circuital, and behavioral levels is essential to fully understand the role of Nav1.2 in neurodevelopmental disorders. However, current mouse models of *SCN2A* only provide limited insight. Canonical *Scn2a* null mice (MGI:2180186, *Scn2a*^*tm1Mml*^) were generated by removing Exon 2 (ΔEx2), which is critical for the function of Nav1.2 [9]. Initial studies of the ΔEx2 *Scn2a* mouse model demonstrated that homozygous (*Scn2a*^−*/*−^) knockout is perinatal lethal, whereas heterozygous (*Scn2a*^*+/*−^) mice are viable and fertile with no major phenotype abnormalities compared to WT mice (*Scn2a*^+/+^) [9].

More recent studies performed with this traditional knock-out mouse model revealed subtle phenotypes. In particular, Dr. Mantegazza’s team in Route des Lucioles, France assessed the behavior of ΔEx2 heterozygous *Scn2a*^*+/*−^mice (male only), they found that ΔEx2 adult *Scn2a*^*+/*−^ male mice have no changes in olfaction or marble burying behaviors [10], similar to our results for HET (*Scn2a*^*gtKO/WT*^) mice. Although they detected some behavioral abnormalities in juveniles, these behavioral abnormalities disappear in adulthood. In contrast, our behavioral tests were all performed on adult mice. Juvenile behavior of *Scn2a*^*gtKO/gtKO*^ mice remains to be tested.

Also, Dr. Yasakawa’s team at RIKEN Center in Japan has conducted extensive behavioral analysis in which most of the assays reveal no difference between WT and *Scn2a*^*+/*−^ (males only) in body weight, hot plate, and auditory startle [13]. Similarly, our study of *Scn2a*^*WT/gtKO*^ males show no statistically significant differences for these tests. However, we noted that female *Scn2a*^*WT/gtKO*^ have a reduced body weight, highlighting the importance of studying both females and males in neuroscience research [33]. Meanwhile Dr. Bender’s group at University of California, San Francisco focused on electrophysiological properties in the ΔEx2 *Scn2a* mice which revealed differences in neuronal excitability and synaptic functions between WT and *Scn2a*^+/−^mice (both males and females); however, no impaired nesting of *Scn2a*^+/−^ mice was observed [11]. Similarly, we do not see any changes in *Scn2a*^*WT/gtKO*^ nesting – only *Scn2a*^*gtKO/gtKO*^ mice display strong impairment in nesting.

A second *Scn2a* null mouse model was achieved by the deletion of exons 4-6 (MGI:6407391, *Scn2a*^*tm1.2Bcgen*^), which was generated and studied by Dr. Kim’s lab at the Korea Advanced Institute of Science and Technology (KAIST) in Daejeon, South Korea. Homozygous *Scn2a* knock-out in this mouse model also died perinatally [12], further illustrating that *Scn2a* null mice cannot be obtained for adult studies. Their phenotypic analysis of heterozygous mice supported the fact that most behaviors of *Scn2a*^*+/−*^ mice are relatively normal compared to WT (males only as well). Both our HET gtKO mice and *Scn2a*^*+/−*^ null mice express ∼50% of *Scn2a* protein expression levels compared to WT mice [34], indicating that a 50% reduction of *Scn2a* protein is not sufficient to induce robust phenotypes in adult mouse behaviors.

Although there are consistent results among these cited studies, inconsistences also exist. Possibly, small differences in handling, maintenance, experimental conditions or genetic background of these mice may contribute to the subtle and variable phenotypes observed in these studies. Genetic background, indeed, has been shown to influence ASD behaviors in mice [35]. By contrast, impaired behaviors (e.g., marble burying and nesting) in *Scn2a*^*gtKO/gtKO*^ mice are clear and severe, highlighting the importance of *Scn2a*^*gtKO/gtKO*^ mice model for the study of *Scn2a*-deficiency from birth to adulthood.

To overcome the existing limitations of current heterozygous *Scn2a* null mouse models, we characterized a gene-trap knockout *Scn2a* mouse model [14], where homozygous animals are viable. Thus, we reasoned that the engineered allele could result in hypomorphic expression of *Scn2a*. The knockout-first strategy has been used to target thousands of genes in mice [14]. Due the fact that the tm1a alleles do not delete exonic sequence, modification of the allele by Cre recombinase is required to produce a null mutation by removal of a critical exon (creating the ‘tm1b’ allele) [14]. The *Scn2a* gtKO involves inserting a lacZ “trapping cassette” into the intron between Exons 1 and 2 and introducing site-specific recombination sites flanking Exons 2-3 of *Scn2a*. We find that the unmodified ‘tm1a’ allele of *Scn2a* (*Scn2a*^*gtKO*^) allows for the expression of residual full-length functional protein, which we believe is the reason why the majority of *Scn2a*^*gtKO/gtKO*^ mice survive to adulthood. In contrast, canonical homozygous knockout (null) of *Scn2a* in mice leads to pups dying 1-2 days after birth with complications including dehydration, hypoxia, ∼16% loss in body weight, and reduced milk content in the stomach [9]. Our *Scn2a*^*gtKO/gtKO*^ mice had a similar ∼15% reduced body weight compared to WT throughout life, but close to 80% of *Scn2a*^*gtKO/gtKO*^ mice survive to adulthood. Another mouse model of neurodevelopmental disorder, *Foxp1*^*-/-*^, also has failure to thrive at weaning unless a soft, high-calorie diet is provided [36] which we similarly provided a gel-based high-calorie food source. The combined phenomenon of failure to thrive in adolescence and overeating in adulthood in humans is also evident in Prader-Willi syndrome [37]. Interestingly, *Scn2a*^*gtKO/gtKO*^ mice seem to have gastrointestinal issues including penile/vaginal and rectal prolapse and diarrhea. *Scn2a* is suggested to be expressed in the enteric nervous system of humans, particularly increased during embryonic weeks 12-16 [38]. In guinea pigs, a study reported that *Scn2a* expressed in the small and large intestines contributes to the somal action potential of enteric sensory neurons [39]. However, whether the gastrointestinal issues of *Scn2a*^*gtKO/gtKO*^ mice is due to the reduced expression of *Scn2a* outside of central nervous system remains to be determined. Another factor to be considered is altered microbiota in *Scn2a*^*gtKO/gtKO*^ mice, as microbiota is suggested to be implication in neurological development [40].

For olfactory acuity discrimination test between water and cinnamon, we found that male *Scn2a*^*gtKO/gtKO*^ mice had a significantly increased interaction time with both scents. However, both male and female *Scn2a*^*gtKO/gtKO*^ mice showed no significant difference from WT in olfactory preference for cinnamon over water. Our olfactory results are similar to a Nav1.2 shRNA knockdown mouse model which also had an increase in time for the mice to discriminate the odorants [41].

Atypical innate behavior (nesting, marble burying, and mating) is an indicator of neurodevelopmental abnormalities in mice. Our *Scn2a*^*gtKO/gtKO*^ mice have little-to-no nest building activity. Decreased nesting has been also been identified in *Mecp2*-mutated mouse model of Rett syndrome [42]. Another common innate phenotypic test performed on mouse models is marble burying, although its translational validity has been under debate [43]. Some studies have suggested marble burying could be a neophobic response [44], repetitive/compulsive [29, 45], or just a normal behavioral routine of inherent burying behavior [46]. Decreased nesting and marble burying were also observed in a Fragile-X mouse model [47]. Since our data reveals that *Scn2a*^*gtKO/gtKO*^ mice bury little to no marbles and hardly touch the nesting material, we suggest that these *Scn2a*^*gtKO/gtKO*^ mice have an impaired neurological development that causes abnormalities of innate behaviors in adulthood.

Mice rely on olfactory sensing for sensory communication [48, 49]. Thus socio-sexual olfactory investigatory preference was used to further understand why the *Scn2a*^*gtKO/gtKO*^ mice display no mating behavior. *Scn2a*^*gtKO/gtKO*^ males show a preference for the male scent instead of the estrus female scent, which was also reported in a Neuroligin-3 mouse model of neurodevelopmental disorders [50]. Future studies are needed to determine the role of *Scn2a* on hormone regulation which is suggested to play a role in neurodevelopment [51, 52]. In addition, *Scn2a* deficiency during development may lead to the up or down regulation of other ion channels or other compensatory mechanisms contributing to the observed phenotypes. This can be studied in future experiments.

In conclusion, the results from this study supports the use of a partial loss-of-function gene-trap allele to model *SCN2A*-deficiency and study genetic sodium channelopathies. Further studies will explore other behavioral, biochemical, and electrophysical properties of these mice at different developmental stages to delineate the differences from WT mice to discern a mechanism of how *Scn2a*-deficiency leads to neurodevelopmental disorders.

## AUTHOR CONTRIBUTIONS

ME, ACP, EL, CMR, and ERC performed behavior and analysis, and husbandry. JZ, ZM, TX, and YL performed the molecular biology and analysis. WS, WK, and YY helped with data analysis and experimental design. YY supervised the project. ME wrote the paper with inputs from YY, WS, ZH, and WK.

### ACKNOWLEDGEMENTS

ME is supported by the National Science Foundation (NSF) Graduate Research Fellowship Program (GRFP) (DGE-1842166). YY is supported by funds from Ralph W. and Grace M. Showalter Research Trust, Purdue Big Idea Challenge 2.0, Purdue Institute for Drug Discovery, and the Purdue Institute for Integrative Neuroscience. We thank Alex Pelle and the Purdue animal staff for their valuable husbandry suggestions.

